# Differential Gene Set Enrichment Analysis: A statistical approach to quantify the relative enrichment of two gene sets

**DOI:** 10.1101/860460

**Authors:** James H. Joly, William E. Lowry, Nicholas A. Graham

## Abstract

Gene Set Enrichment Analysis (GSEA) is an algorithm widely used to identify statistically enriched gene sets in transcriptomic data. However, to our knowledge, there exists no method for examining the enrichment of two gene sets relative to one another. Here, we present Differential Gene Set Enrichment Analysis (DGSEA), an adaptation of GSEA that assesses the relative enrichment of two gene sets. Using the metabolic pathways glycolysis and oxidative phosphorylation as an example, we demonstrate that DGSEA accurately captures the hypoxia-induced shift towards glycolysis. We also show that DGSEA is more predictive than GSEA of the metabolic state of cancer cell lines, including lactate secretion and intracellular concentrations of lactate and AMP. Furthermore, we demonstrate that DGSEA identifies novel metabolic dependencies not found by GSEA in cancer cell lines. Together, these data demonstrate that DGSEA is a novel tool to examine the relative enrichment of two gene sets.

## Introduction

Given the ever-increasing availability of -omics data characterizing the genome, transcriptome, proteome, and metabolome, there is a persistent need for approaches that extract biological insights from these complex data sets. Gene Set Enrichment Analysis (GSEA) (1) has proved to be one of the most popular and powerful tools for analyzing transcriptomic data. In addition, the GSEA algorithm has proved useful for the analysis of other data types including DNA copy number alterations (2), proteomics (3, 4), phospho-proteomics (5, 6), and metabolomics (7, 8). Regardless of the data type, the key concept underlying GSEA is that pre-defined sets of functionally related genes can display significant enrichment that would be missed by examination of individual genes. By using the entire dataset as background, researchers can identify pathways upregulated and downregulated in phenotype(s) of interest. The GSEA approach is widely successful and has inspired many extensions, improvements, and variations to analyze individual gene sets (9–14).

However, to our knowledge, no tools exist for analyzing the difference between two gene sets. Because biological control involves many tradeoffs and pathway branches, statistical methods are needed to accurately measure how two gene sets or pathways are coordinately regulated with respect to each other. Here, we present Differential Gene Set Enrichment Analysis (DGSEA), an adaption of GSEA that calculates the enrichment of two pathways relative to each other. Using the metabolic pathways glycolysis and oxidative phosphorylation as a test case, we demonstrate the application of DGSEA to gene expression data accurately captures cellular phenotypes. First, testing DGSEA with gene expression data from a panel of cell lines subjected to hypoxia, we found that DGSEA accurately identified the hypoxia-induced tradeoff between glycolysis and oxidative phosphorylation. Next, by applying DGSEA to two panels of cancer cell lines with paired gene expression and metabolomic data, we demonstrated that DGSEA is more predictive of cellular metabolism than GSEA, specifically lactate secretion rates and intracellular concentrations of lactate and AMP. Finally, through integration of DGSEA with genome-wide genetic dependency screens, we found that DGSEA identified novel metabolic dependencies that were missed by GSEA in cancer cell lines. As the availability of -omics data continues to increase, DGSEA will serve as a tool to identify tradeoffs between gene sets or pathways that govern biological control.

## Results

### DGSEA quantifies the enrichment of two gene sets relative to each other

To quantify the enrichment of two gene sets relative to each other, we adapted GSEA to create Differential Gene Set Enrichment Analysis (DGSEA). Given two gene sets of interest (e.g., Gene Sets A and B), the goal of DGSEA is to determine whether the members of Gene Sets A and B are randomly distributed with reference to each other. If Gene Sets A and B are upregulated and downregulated, respectively, we expect that A and B will be at opposite sides of the rank list. Although we use the terminology “gene set”, the DGSEA algorithm is agnostic to data type and can be used with genomic, transcriptomic, proteomic, or metabolomic data alongside sets of functionally related genes, proteins, or metabolites. DGSEA first ranks the data by any suitable metric including fold change, signed p-value, signal-to-noise ratio, or correlation with a phenotype of interest (Fig. 1). Second, we calculate an enrichment score (ES) for each gene set (ES_A_ and ES_B_) by walking down the rank list and finding the maximum deviation from zero of a running-sum, weighted Kolmogorov-Smirnov-like statistic. This is equivalent to the GSEA algorithm. Then, the difference between ES_A_ and ES_B_ is calculated to measure the enrichment of the two gene sets relative to each other (ES_AB_ = ES_A_ - ES_B_). We then estimate the significance level of ES_A_, ES_B_, and ES_AB_ using an empirical permutation test as in the original GSEA algorithm. Specifically, the rank list is permuted, permutation enrichment scores are calculated for each gene set (pES_A_, pES_B_, and pES_AB_), and then the nominal p-value is calculated relative to the same-signed portion of the null distribution. Next, the normalized enrichment score (NES) is calculated by dividing positive and negative ES by the mean of positive or negative pES, respectively. Finally, to estimate the false discovery rate (FDR), a null distribution of NES values is generated using a list of background gene sets. Background gene sets can be chosen based on the biological meaning of the tested gene sets, or they can be randomly generated. Using the background gene sets, the null distribution is the union of NES values comparing Gene Set A versus all background pathways (NES_AX_) with the NES values comparing all background gene sets versus Gene Set B (NES_XB_). The combination of these distributions is termed NES_XY_. For a given NES_AB_, the FDR is then calculated as the ratio of the percentage of the same-signed NES_XY_ greater than or equal to NES_AB_ divided by the percentage of NES_XY_ with the same sign as NES_AB_. The FDR estimates for NES_A_ and NES_B_ are generated using a similar approach based on single background gene sets, equivalent to GSEA. The output of DGSEA is thus the relative enrichment and statistical significance of Gene Set A versus B, as well as the individual enrichment and statistical significance of Gene Sets A and B.

**Figure 1.**
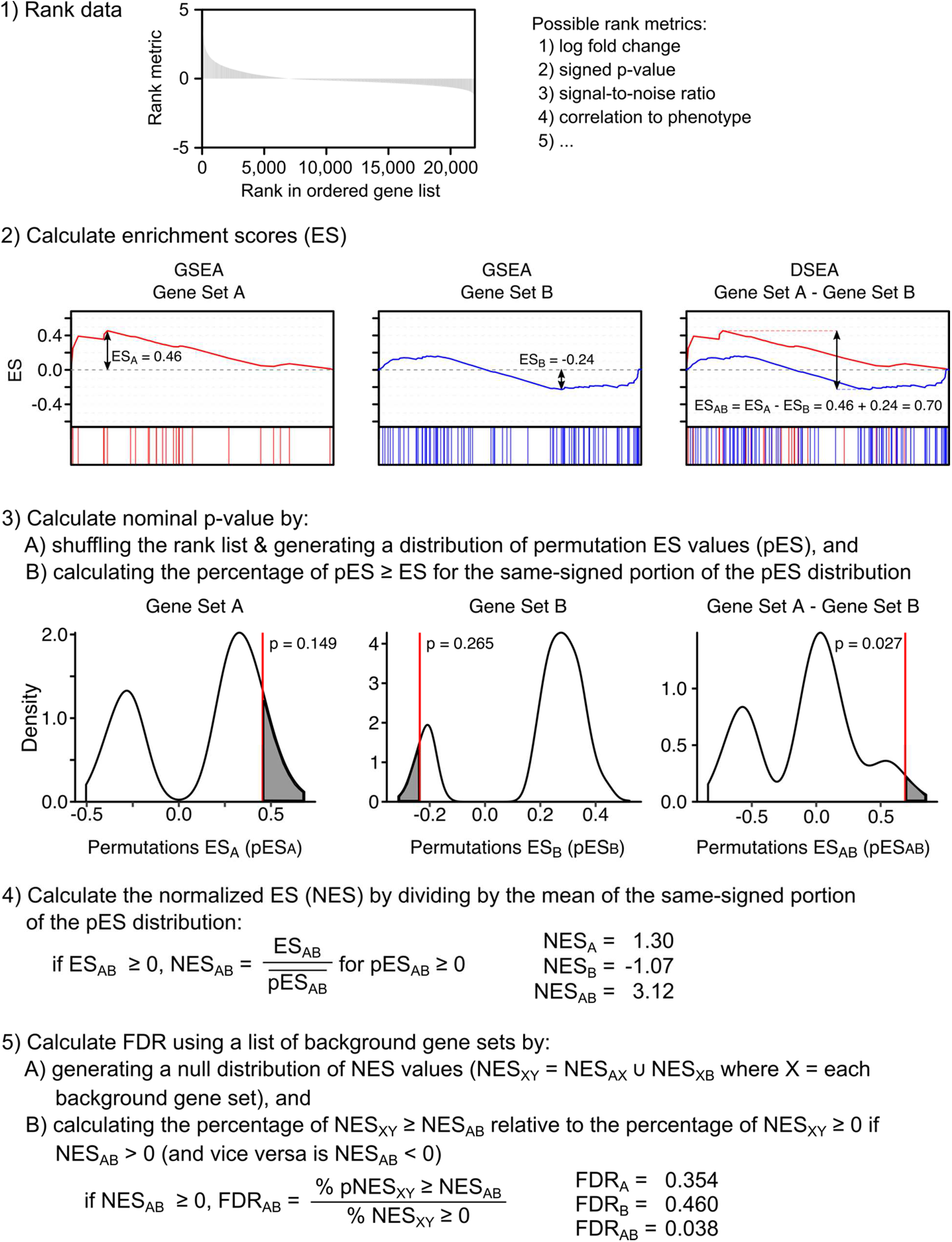
Differential Gene Set Enrichment Analysis (DGSEA) quantifies the enrichment between two gene sets relative to each other. A data set is first ranked by any suitable metric inducing fold change, signed p-value, signal-to-noise ratio, or correlation with phenotype. Second, the enrichment score (ES) is calculated for each individual gene set by walking down the rank list and finding the maximum deviation from zero of a running-sum, weighted Kolmogorov-Smirnov-like statistic (ES_A_ and ES_B_, equivalent to GSEA, left & middle). Then, the difference between the enrichment scores of two gene sets is calculated (ES_AB_= ES_A_ - ES_B_, right). Third, the statistical significance of ES_A_, ES_B_, and ES_AB_ is estimated by using an empirical permutation test that preserves the structure of the original data. Specifically, a null distribution is generated by shuffling the rank list and calculating the permutation ES values (e.g., pES_A_), and the nominal p-value is calculated relative to the same-signed distribution. Fourth, the normalized enrichment score (NES) is then calculated by dividing the observed ES by the mean of the same-signed portion of the permutation ES distribution 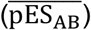. Fifth, to control for false discovery rate (FDR), a null distribution of NES values is generated using a list of background gene sets. For DGSEA, the null distribution of NES values (NES_XY_) is the union of the NES values comparing Gene Set A versus each background gene set X (NES_AX_) and the NES values comparing each background gene set X against Gene Set B (NES_XB_). For a given NES_AB_ greater than zero, the FDR is then calculated as the ratio of the percentage of all NES_XY_ greater than or equal to NES_AB_ divided by the percentage of observed NES_XY_ is positive and similarly if NES_AB_ is negative.

### DGSEA accurately captures the coordinate upregulation of glycolysis and downregulation of oxidative phosphorylation in hypoxia

Hypoxia is associated with a metabolic shift away from oxidative phosphorylation and towards glycolysis. In the absence of oxygen, hypoxia-inducible factor 1-α (HIF1α) transcriptionally activates glucose catabolism through expression of glucose transporters, glycolytic enzymes, and lactate dehydrogenase A (15). Moreover, the HIF1α-target gene pyruvate dehydrogenase kinase 1 (*PDK1*) suppresses metabolic flux from pyruvate to acetyl-CoA, diverting carbon away from the mitochondria and thereby reducing oxidative phosphorylation (16). We therefore first tested the ability of DGSEA to detect the hypoxic shift from oxidative phosphorylation to glycolysis. We applied DGSEA to RNASeq data from 31 breast cancer cell lines subjected to either 1% or 20% oxygen (17) either as paired cell lines (i.e., hypoxia / normoxia) or as individual samples (i.e., single sample DGSEA) (Fig. 2A). Analysis with a consensus hypoxia gene set confirmed that all 31 cell lines demonstrated enrichment of hypoxia-regulated genes upon exposure to 1% oxygen (Supplemental Fig. 1A). To test the differential enrichment of glycolysis and oxidative phosphorylation, we chose gene sets A) core glycolysis (hsaM00001, which includes conversion of glucose into pyruvate) and B) oxidative phosphorylation (hsa00190), respectively, from KEGG (18). For the paired cell line analysis, DGSEA was significantly upregulated in 21 of 31 cell lines (p-value < 0.05, FDR < 0.25, Fig. 2B and Supp. Table 1). For the individual pathways, GSEA demonstrated significant upregulation of core glycolysis in all 31 cell lines but significant downregulation of oxidative phosphorylation in only 21 of 31 cell lines. Surprisingly, upon exposure to 1% oxygen, 10 cell lines exhibited significant <u>*upregulation*</u> of oxidative phosphorylation. One cell line, MCF12A, had a similar induction of both core glycolysis and oxidative phosphorylation in 1% oxygen (Fig. 2C). Notably, the cell lines with upregulated oxidative phosphorylation in 1% oxygen were the same cell lines identified by DGSEA as not having a significantly differential response between core glycolysis and oxidative phosphorylation. Substitution of the full glycolysis pathway (Glycolysis-Gluconeogenesis, hsa00010) as Gene Set A or TCA cycle (hsa00020) as Gene Set B yielded very similar results to usage of Core Glycolysis (hsaM00001) and Oxidative Phosphorylation (hsa00190) as Gene Sets A and B, respectively (not shown).

**Figure 2.**
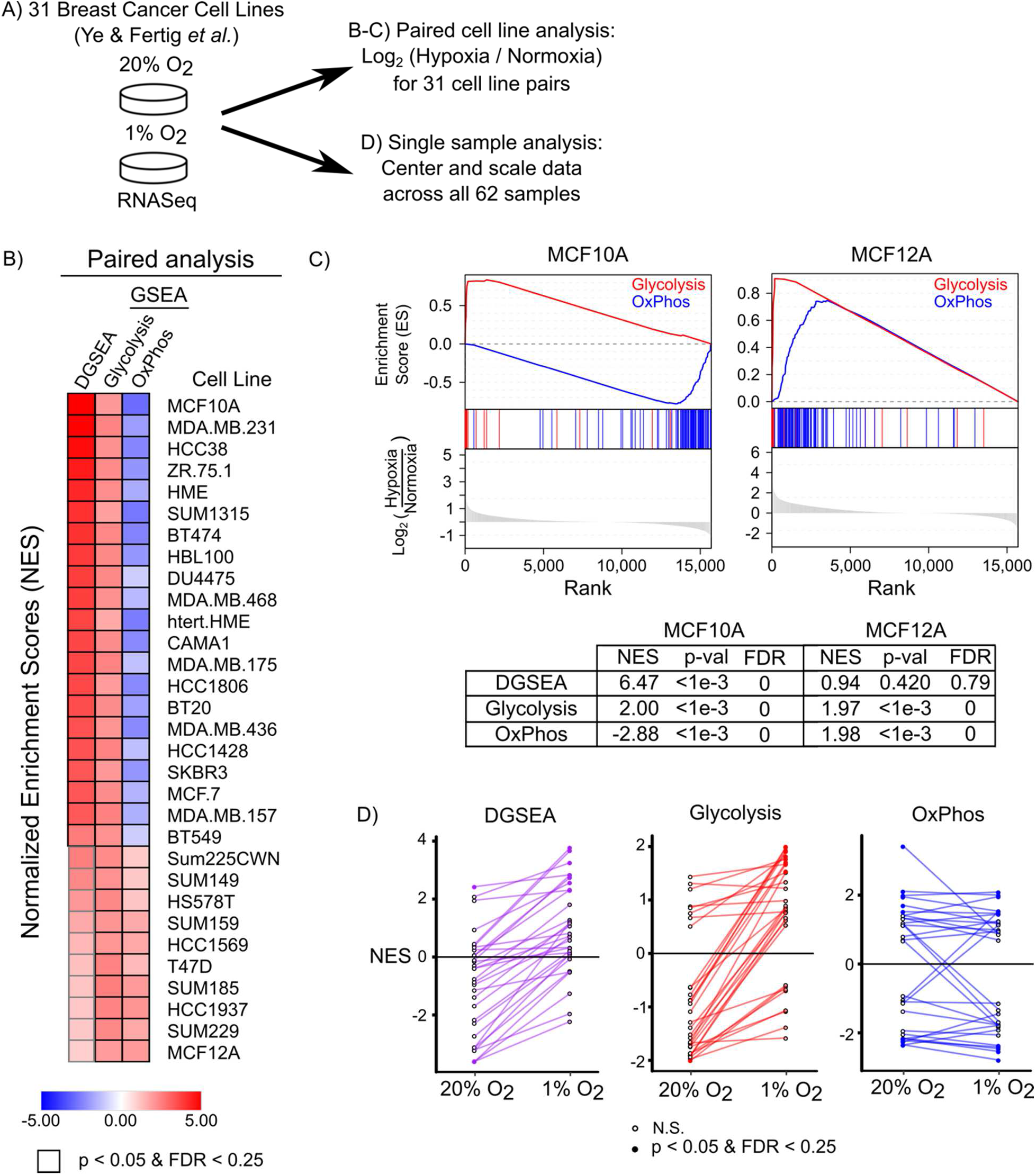
DGSEA accurately captures the coordinate upregulation of glycolysis and downregulation of oxidative phosphorylation in hypoxia. **A)** Schematic of data normalization methods used to generate gene ranking metrics using RNAseq data from 31 breast cancer cell lines subjected to hypoxia (1% O_2_) or normoxia (20% O_2_) (17). Genes were first ranked by the paired fold-change of hypoxia over normoxia for analysis in panels B and C. For panel D, gene expression levels from 62 individual samples were centered, scaled and then analyzed by single sample DGSEA (ssDGSEA) or ssGSEA. **B)** DGSEA on paired cell line data identified cell lines with coordinately increased glycolysis and decreased oxidative phosphorylation. DGSEA (Core Glycolysis-OxPhos) and GSEA (Core Glycolysis & OxPhos) Normalized Enrichment Scores (NES) were calculated for each cell line. Some, but not all, cancer cell lines increased Glycolysis and decreased OxPhos when subjected to hypoxia. Black outline denotes p-value < 0.05 and FDR < 0.25. **C)** Representative mountain plots and table of values for two cell lines subjected to hypoxia. MCF10A cells exhibited strong upregulation of glycolysis and downregulation of OxPhos, whereas MCF12A cells exhibited a strong induction of both glycolysis and OxPhos. **D)** Line plots for ssDGSEA and ssGSEA for each cell line show the change in NES upon switch from 20% to 1% oxygen. DGSEA and glycolysis were upregulated in most cell lines, whereas OxPhos showed neutral changes in most cell lines. Accordingly, DGSEA showed small increases that reflect increased glycolysis but not decreased OxPhos. Filled circles denote nominal p-value < 0.05 and FDR < 0.25. Open circles denote not significant (N.S.).

We next tested whether similar trends were observable in single sample DGSEA (ssDGSEA) and GSEA (ssGSEA). Again, analysis with a consensus hypoxia gene signature confirmed that cell lines responded to 1% oxygen (Supp. Fig. 1B). Using DGSEA to compare the relative enrichment of core glycolysis and oxidative phosphorylation, we found that nearly every cell line increased its NES score in 1% oxygen (Fig. 2D). In addition, the number of cell lines with significant ssDGSEA scores at 1% oxygen was increased relative to 20% oxygen (7 versus 1). Consistent with the paired cell line analysis, most cell lines increased the ssGSEA NES of core glycolysis in 1% oxygen. For oxidative phosphorylation, however, most cell lines had only slightly negative or negligible changes in the ssGSEA NES between 20% and 1% oxygen. Taken together, these results demonstrate although not all cell lines subjected to hypoxia exhibit gene expression signatures consistent with shifting from oxidative phosphorylation to glycolysis, DGSEA does accurately identify cell lines which exhibit the hypoxia-induced coordinate upregulation and downregulation of glycolysis and oxidative phosphorylation, respectively.

### DGSEA is more predictive than GSEA of lactate secretion and glucose consumption in cancer cell lines

Lactate secretion is often used as a marker of the metabolic shift between glycolysis and oxidative phosphorylation. Because DGSEA can measure the relative difference between glycolysis and oxidative phosphorylation, we therefore hypothesized that DGSEA would be more predictive of lactate secretion rates than GSEA using either the glycolysis or oxidative phosphorylation gene sets alone. To test this hypothesis, we analyzed paired gene expression and metabolite consumption and secretion rates from the NCI-60 panel of cancer cell lines (19) (Fig. 3A and Supp. Table 2). Indeed, we found that ssDGSEA NESs were more significantly correlated with lactate secretion rates than either core glycolysis or oxidative phosphorylation ssGSEA NESs (Fig. 3B). Similar to our results with hypoxia, we found that ssDGSEA was a better predictor of lactate secretion than ssGSEA for all combinations of similar gene sets (e.g., Gene Set A is either Core Glycolysis or Glycolysis-Gluconeogenesis and Gene Set B is either Oxidative Phosphorylation or TCA Cycle) (Supp. Fig. 2). Interestingly, ssDGSEA was also more significantly predictive of glucose uptake across the 59 cell lines than ssGSEA using either glycolysis or oxidative phosphorylation. Together, these results reveal that DGSEA was more predictive of lactate secretion and glucose consumption than GSEA across a panel of cancer cell lines.

**Figure 3.**
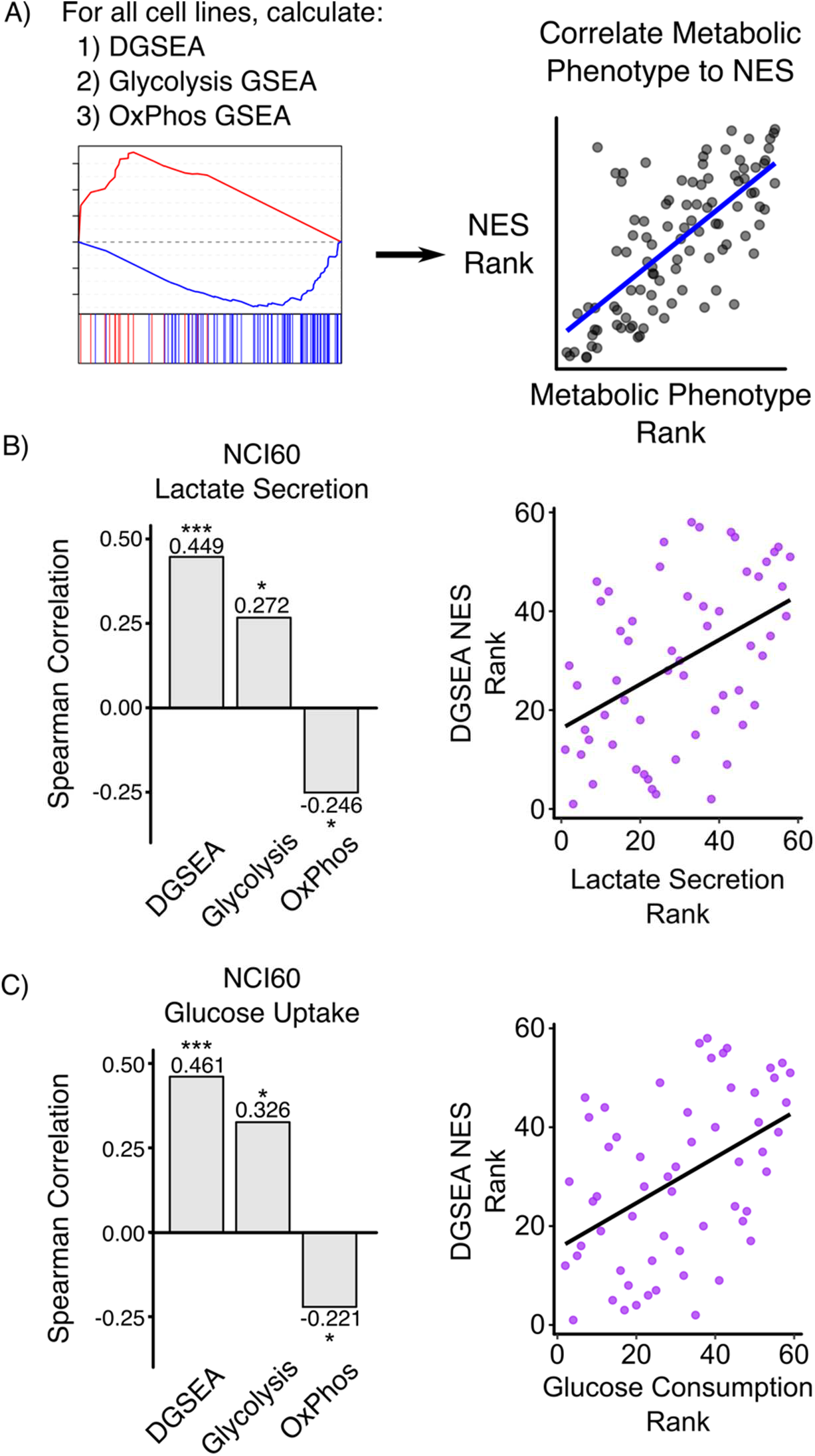
DGSEA is a better predictor of cellular metabolism than GSEA. **A)** Schematic of process used to correlate pathway activity, as measured by GSEA or DGSEA, with metabolic phenotypes. **B-C)** DGSEA more accurately predicted lactate secretion and glucose uptake rates than GSEA. Gene expression data was centered and scaled across 59 of the NCI-60 cancer cell lines and the NES values for Core Glycolysis, OxPhos, and DGSEA were calculated for each cell line. Spearman rank correlation coefficients were calculated between each NES and lactate secretion or glucose uptake data (19). Scatter plots showing the spearman correlation are shown (right). * indicates p < 0.05, ** p < 0.01, and *** p < 0.001.

### DGSEA is correlated with high concentrations of intracellular lactate concentrations and low concentrations of intracellular AMP in adherent cell lines

Next, we tested how DGSEA correlated with intracellular metabolite concentrations. Although intracellular metabolite concentrations do not reflect pathway flux, we hypothesized that comparing DGSEA and steady-state metabolite abundance would reveal trends consistent with coordinate upregulation of glycolysis and downregulation of oxidative phosphorylation. For this purpose, we used paired RNAseq and metabolomics data from 897 cancer cell lines from the Cancer Cell Line Encyclopedia (20). Since culture type has been reported to be a major determinant of metabolism, we separately analyzed cancer cell lines cultured in adherent and suspension cultures. Correlating DGSEA NESs for 836 adherent cell lines against 225 intracellular metabolite concentrations, we found that the metabolite most correlated with DGSEA was 1-methylnicotinamide (MNA), which has no known role in regulation of glycolysis or oxidative phosphorylation (Fig. 4A and Supp. Table 3). However, the second most correlated metabolite with DGSEA NES was lactate, suggesting that DGSEA accurately captured the tradeoff between glycolysis and oxidative phosphorylation. As with the lactate secretion data (Fig. 3), we found that DGSEA NESs correlated better with intracellular lactate levels than did GSEA NESs using either glycolysis or oxidative phosphorylation alone (Fig. 4B and Supp. Fig. 3A). Interestingly, we found that the metabolite most anticorrelated with DGSEA NESs was AMP, one of the classical readouts of the cellular energy state (21). DGSEA again was a better predictor of AMP levels than GSEA with either glycolysis of oxidative phosphorylation alone. Notably, these results with adherent cultures were not recapitulated in suspension cultures, perhaps due to sample size limitations (Supp. Fig. 3B). Taken together, these results indicate that DGSEA testing the relative enrichment between glycolysis and oxidative phosphorylation strongly correlated with steady state levels of metabolites that indicate the tradeoff between glycolysis and oxidative phosphorylation (i.e., lactate) and anticorrelated with metabolites indicative of a low energetic state (i.e., AMP).

**Figure 4.**
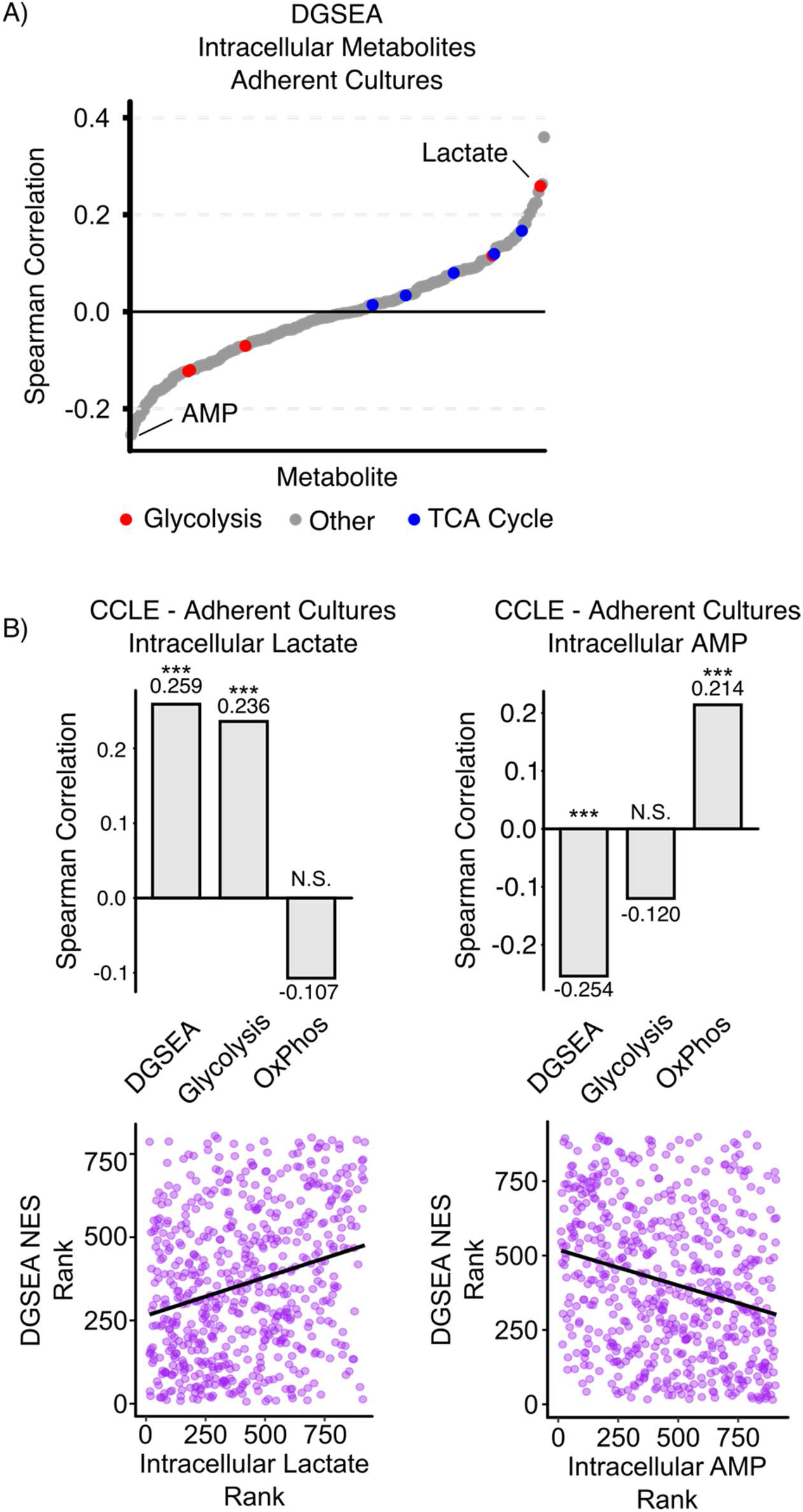
DGSEA is a better predictor of intracellular lactate and AMP levels than GSEA for adherent cell cultures. **A)** Increased intracellular lactate and decreased AMP correlated with increased glycolysis and decreased OxPhos. RNASeq data was centered and scaled across all adherent cell culture lines in the Cancer Cell Line Encyclopedia and then the spearman correlation coefficient was calculated between DGSEA NESs and metabolite abundances. Lactate was the second most correlated metabolite and AMP was the least correlated metabolic with DGSEA. **B)** Barplots showing the comparison of DGSEA and glycolysis and OxPhos GSEA. Scatter plots showing the correlation between DGSEA and lactate or AMP are shown. *** indicates p < 0.001.

### DGSEA provides novel insight into the metabolic dependencies of cancer cell lines

Recent developments in gene essentiality screening have enabled the identification of novel gene dependencies in cancer cell lines (22–25). Because DGSEA can identify the metabolic tradeoff between glycolysis and oxidative phosphorylation, we hypothesized that DGSEA would identify different essential genes than GSEA using either glycolysis or oxidative phosphorylation alone. To test this hypothesis, we first calculated NESs for DGSEA comparing glycolysis and oxidative phosphorylation and GSEA for glycolysis and oxidative phosphorylation across 1,019 cell lines in the Cancer Cell Line Encyclopedia (Fig. 5A). Then, for each gene in the Cancer Dependency Map (DepMap), we calculated the correlation coefficient between cell line NESs and the gene dependency score (CERES score) and ranked genes by correlation coefficient (23). Examination of DGSEA rank list revealed that the top-ranked gene was mitochondrial aconitase (*ACO2*) and that six of the top 10 genes were members of the oxidative phosphorylation pathway (*ATP5F1*, *NDUFA6*, *ATP5H*, *ATP5I*, *ATP5O*, and *COX10*) (Fig. 5B and Supp. Table 4). Notably, the essentiality of the F-Type ATPase genes (mitochondrial complex V) was increased in DGSEA relative to GSEA using either glycolysis or oxidative phosphorylation alone (Fig. 5C). Among the genes most anticorrelated with DGSEA was the transcription factor *JUN*. For GSEA using glycolysis alone, the gene whose CERES score most correlated with NES was PIK3CA (Supp. Fig. 4). *ACO2* was the second most correlated gene with glycolysis NES, but the correlation was weaker than with DGSEA NES. Taken together, these results demonstrate that DGSEA reveals unique genetic vulnerabilities in cancer cells with coordinately upregulated glycolysis and downregulated oxidative phosphorylation.

**Figure 5.**
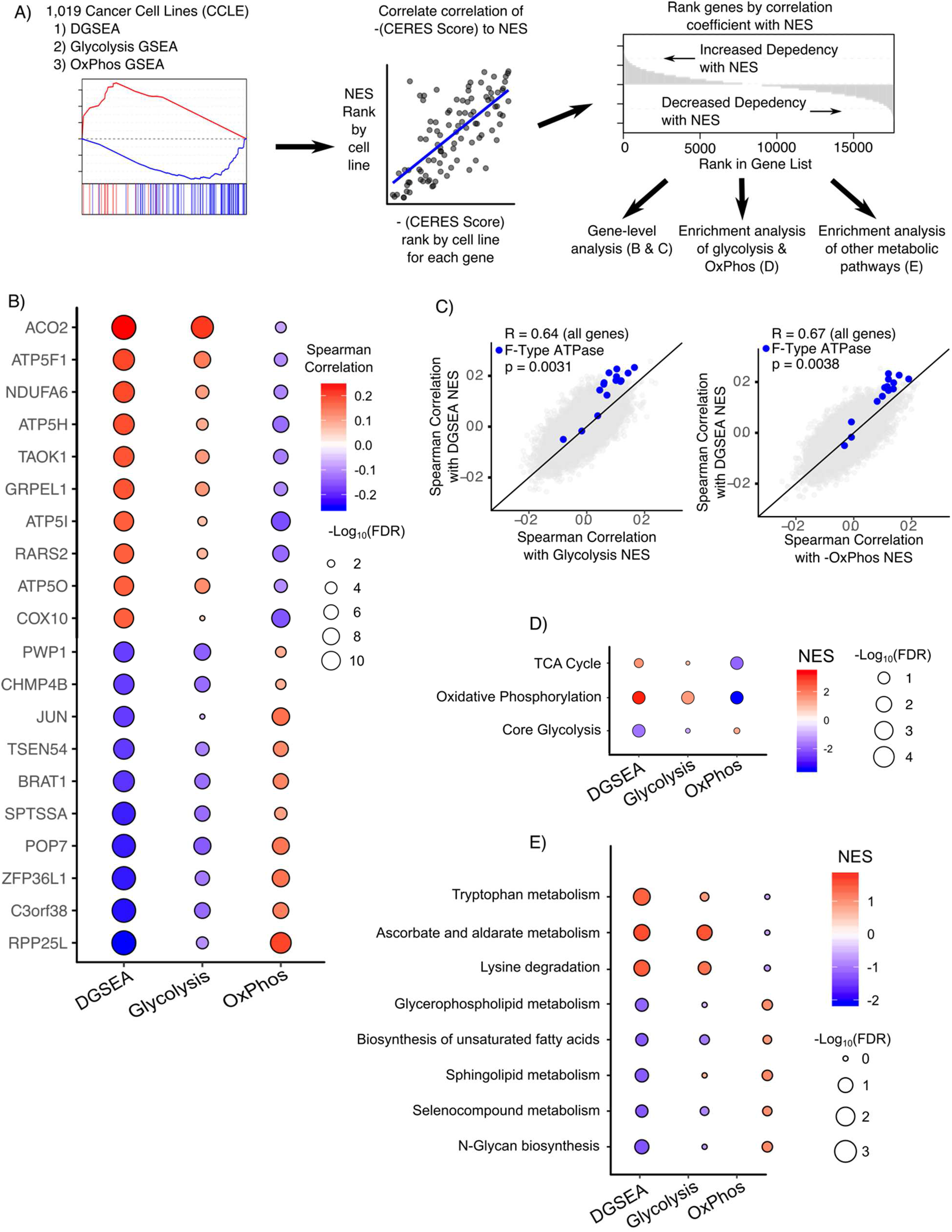
DGSEA reveals novel metabolic dependencies. **A)** Schematic outlining workflow used to identify metabolic dependencies with DGSEA or GSEA using glycolysis and oxidative phosphorylation. **B)** DGSEA reveals novel genetic dependencies compared to GSEA. RNAseq data was scaled and centered for 1,019 cancer cell lines and the DGSEA or GSEA NES was calculated for each cell line. Spearman correlations were calculated between dependency (CERES score) and the normalized enrichment scores (NES) from DGSEA or GSEA. Spearman correlations for the top 10 and bottom 10 most correlated genes with DGSEA are shown. The circle color indicates the correlation coefficient and the circle size indicates the FDR-corrected p-value. **C)** DGSEA exhibits increased dependency on F-Type ATPase genes compared to glycolysis and OxPhos. Since positive DGSEA reflects a negative OxPhos value, each OxPhos NES was multiplied by negative one. Reference lines for y=x are drawn in black. P-values were calculated for enrichment based on the cumulative distribution function of the hypergeometric distribution. **D-E)** RNAseq data was scaled and centered for 1,019 cancer cell lines and DGSEA or GSEA NES values were calculated for each cell line. Spearman correlations were calculated between dependency (CERES score) and the normalized enrichment scores (NES) from DGSEA or GSEA. Then, GSEA was run against all KEGG metabolic pathways to identify pathways that exhibit increased or decreased dependency with DGSEA.

Next, we tested whether the DGSEA and GSEA rank lists were enriched for the metabolic pathways used to generate the DGSEA and GSEA NESs, namely glycolysis and oxidative phosphorylation. As expected based on the enrichment of individual oxidative phosphorylation genes in the DGSEA rank list, sensitivity to depletion of the oxidative phosphorylation pathway was enriched in cell lines with large positive DGSEA NESs (Fig. 5D and Supp. Table 5). Surprisingly, we found that the DGSEA rank list was strongly de-enriched for glycolysis genes. That is, cell lines with large positive DGSEA NESs (i.e., relative upregulation of glycolysis and downregulation of oxidative phosphorylation) were relatively insensitive to depletion of glycolytic genes. In contrast, the essentiality of glycolytic genes was increased in cell lines with large negative DGSEA NESs (i.e., relative downregulation of glycolysis and upregulation of oxidative phosphorylation). Interestingly, this trend was not replicated in either rank lists for GSEA with either glycolysis or oxidative phosphorylation. Examining the rank list for GSEA with oxidative phosphorylation, we found that increased oxidative phosphorylation correlated with a decreased dependence on genes in the oxidative phosphorylation and TCA Cycle gene sets. However, the same finding did not hold true for glycolysis. Taken together, these results suggest that gene essentiality for glycolysis increases only when glycolysis is low and oxidative phosphorylation is high (i.e, large negative DGSEA NES).

We next sought to understand which metabolic pathway(s) other than glycolysis and oxidative phosphorylation exhibited changes in essentiality when glycolysis and oxidative phosphorylation are coordinately upregulated and downregulated, respectively (i.e., large positive DGSEA NESs). Interestingly, cell lines with large DGSEA NES exhibited increased dependence upon the Tryptophan Metabolism pathway (Fig. 5E and Supp. Fig. 5). In contrast, neither GSEA with glycolysis nor with oxidative phosphorylation alone exhibited increased dependency on Tryptophan Metabolism. In addition, the pathways Ascorbate and Aldarate Metabolism (hsa00053) and Lysine Degradation (hsa00310) also exhibited increased dependency with DGSEA. We next asked which metabolic pathways exhibit increased dependency upon a shift towards oxidative phosphorylation and away from glycolysis (i.e., large negative DGSEA NES). In addition to Core Glycolysis, we found that cell lines exhibited increased dependence on N-Glycan Biosynthesis (hsa01100), Sphingolipid Metabolism (hsa00600), and Biosynthesis of Unsaturated Fatty Acids (hsa01040) (Fig. 5E).

## Discussion

Traditional gene set enrichment analyses are limited to examining one set of genes at a time. Our DGSEA method builds upon the original GSEA algorithm to measure the enrichment of two gene sets relative to each other. To our knowledge, DGSEA is the first statistical framework to directly quantify the relative enrichment of two gene sets. DGSEA can be run using traditional ranking metrics (e.g., fold-change between perturbation and control) or using single-sample methods across many samples (i.e., ssDGSEA). In this way, DGSEA provides similar usability to GSEA while serving as an extension to pathway analysis not currently possible in other approaches. Our DGSEA software is freely available at https://github.com/JamesJoly/DGSEA.

To test the accuracy of DGSEA, we first used hypoxia as an example of a metabolic shift between glycolysis and oxidative phosphorylation. Indeed, we found that DGSEA accurately captured the metabolic tradeoff between upregulated glycolysis and downregulated oxidative phosphorylation (Fig. 2). Notably, DGSEA identified a metabolic switch in only 21 of 31 cell lines, a finding confirmed by the observation that the 10 other cell lines increased oxidative phosphorylation in response to hypoxia. These surprising findings may be explained by the fact that some cell lines require concentrations of oxygen lower than 1% to suppress oxidative phosphorylation (26). Another explanation is that hypoxia gene signatures correlate with signatures of both glycolysis and oxidative phosphorylation in single-cell RNAseq data (27). Regardless, in cell lines with the classic hypoxia-induced metabolic shift, DGSEA correctly identified coordinate increases in glycolysis and decreases in oxidative phosphorylation.

Having established the accuracy of DGSEA, we proceeded to analyze how DGSEA using the glycolysis and oxidative phosphorylation pathways correlated with traditional metrics of cellular metabolism, namely lactate secretion and glucose consumption (Fig. 3). Our finding that DGSEA more accurately predicted lactate secretion rates than either GSEA with glycolysis or oxidative phosphorylation alone confirmed that DGSEA accurately captured the tradeoff between upregulated glycolysis and downregulated oxidative phosphorylation. In addition, the fact that DGSEA also was more predictive of glucose consumption than GSEA may reflect the highly glycolytic nature of these cancer cell lines in tissue culture. Furthermore, we found that DGSEA NESs of adherent cancer cell lines were more correlated with intracellular lactate than either GSEA with glycolysis or oxidative phosphorylation (Fig. 4). Although steady-state levels of lactate do not necessarily reflect the relative activity of glycolysis and oxidative phosphorylation, they do suggest that DGSEA reflects the balance between conversion of pyruvate to lactate and acetyl-CoA for the TCA cycle. In addition, we found that DGSEA NESs were more significantly anticorrelated with intracellular levels of AMP than either GSEA with glycolysis or oxidative phosphorylation alone. Notably, AMP regulates both glycolysis and oxidative phosphorylation through AMP-activated kinase (AMPK). When the ratio of AMP to ATP is high, AMPK stimulates glycolysis through phosphorylation of glycolytic enzymes and also upregulates mitochondrial biogenesis (21). Because the analyzed metabolomic data did not include ATP levels, we cannot calculate the AMP:ATP ratio to infer the activity of AMPK in these cell lines. However, taken together these results demonstrate that DGSEA analysis is highly informative for metabolic pathway activity and intracellular energetic state.

We also found that applying DGSEA to glycolysis and oxidative phosphorylation gave novel insights into metabolic dependencies. The genes that exhibited increased dependency when glycolysis is high and OxPhos is low (i.e., large positive correlation with DGSEA NES) included many genes comprising mitochondrial complex V, the ATP synthase involved in oxidative phosphorylation in the mitochondria. Indeed, DGSEA was a better predictor of sensitivity to depletion of oxidative phosphorylation genes than either GSEA with glycolysis or oxidative phosphorylation alone (Fig. B-D). The increased essentiality for Complex V when glycolysis is upregulated relative to OxPhos is particularly interesting given that high DGSEA indicates increased conversion of glucose to lactate, which would produce less ATP than OxPhos. As such, these results imply that ATP production from Complex V becomes more essential when carbon is diverted away from OxPhos. These results are notable because even highly glycolytic cancer cells require mitochondrial metabolism for survival (28). The most effective to date cancer drugs targeting mitochondrial metabolism are the biguanides metformin and phenformin, which inhibit mitochondrial complex I (29, 30), but our results suggest that inhibitors of mitochondrial complex V might be effective in tumor cells with high glycolysis and low oxidative phosphorylation. Interestingly, whereas DGSEA identified sensitivity to depletion of oxidative phosphorylation genes, GSEA using the glycolysis pathway alone identified that *PIK3CA* was the gene most correlated with increased glycolysis NESs. This result is consistent with the observation that PI3K-AKT signaling stimulates glycolysis (31) and can render cancer cells dependent on glucose for survival (32).

Our dependency analysis on the metabolic pathway-level revealed an unexpected dependency on tryptophan metabolism in cancer cells with high glycolysis and low oxidative phosphorylation (i.e., high DGSEA NES). This tryptophan result is interesting in light of the fact that catabolism of tryptophan to produce kynurenine is associated with immunosuppression (33). Tryptophan catabolism may also have beneficial roles for tumor progression independent of the immune response, including driving tumor growth in an autocrine fashion (34). Our results suggest that further investigation into how the upregulation of glycolysis and downregulation of oxidative phosphorylation may affect the efficacy of enzymatic inhibitors of tryptophan catabolism, including indoleamine-2,3-dioxygenase (IDO) and tryptophan-2,3-dioxygenase (TDO), which are currently in clinical trials (35). In contrast, when glycolysis is low and OxPhos is high (i.e., low DGSEA NES), we found increased dependence on many lipid metabolism pathways (Fig. 5E). The link to increased dependence on lipid metabolic pathways is notable given that reductive glutamine metabolism has been shown to support lipid synthesis under hypoxia (36, 37). Additionally, increased dependence on fatty acid biosynthesis may reflect a cellular need to upregulate alternative carbon sources to fuel oxidative phosphorylation when glycolytic flux is low. Together, these results demonstrate that applying DGSEA in concert with gene essentiality screening enables the identification of novel genes and pathways essential for cancer cell survival.

In summary, DGSEA is a novel framework for analyzing the tradeoffs between two gene sets or pathway. Although we applied DGSEA to glycolysis and oxidative phosphorylation as a proof-of-concept, we believe that the DGSEA framework is generalizable to examine other gene sets which may exhibit coordinate regulation. Furthermore, since GSEA has been demonstrated to work on other -omic layers, we speculate that DGSEA will accurately capture trade-offs in phospho-proteomic and metabolomic data. As our understanding of biological control becomes more advanced, DGSEA will serve as a useful tool to accurately reflect cellular phenotypes.

## Methods

### Enrichment Score ES_S_ & ES_AB_ Calculation

The enrichment score ES is calculated using the Gene Set Enrichment Analysis (1) algorithm:

1. Rank the order of N genes in expression dataset according to any suitable metric inducing fold change, signed p-value, signal-to-noise ratio, or correlation with phenotype.
2. Evaluate the fraction of genes in set S weighted by their correlation with the phenotype and the fraction of genes not in S.
3. The Enrichment Score *ES*_*S*_ is the maximum deviation from zero of P_hit_ – P_miss_. The score is calculated by walking down the gene list L and increasing a running-sum statistic by P_hit_ when encountering a gene in S and decreasing it by P_miss_ when encountering a gene not in S.
4. For calculating *ES*_*AB*_ comparing two gene sets, the maximum deviation from zero for gene set B is subtracted from the maximum deviation from zero for gene set A.

### Estimating Significance

We estimate significance of the ES by using an empirical permutation test procedure that preserves the ranking metric. For each permutation, we shuffle the gene labels and recompute the ES of the gene set for the shuffled data (pES) to generates a null distribution of pES. The p-value is then calculated relative to this null distribution, using the positive or negative portion of the distribution corresponding to the sign of the observed ES (Fig. 1).

### Adjustment for Multiple Hypothesis Testing

We adjust the estimated significance level for differences for ES_AB_ by generating a null distribution that compares A & B separately to all non-A&B (background) gene sets, hence referred to as ES_XY_. We then normalize each ES for each gene set to account for differences in gene set size, separately rescaling the positive and negative scores by dividing by the mean of the same-signed portion of the pES distribution. Finally, for a given NES_AB_ ≥ 0, the FDR is calculated as the ratio of the percent for all NES_XY_ ≥ NES_AB_ divided by the percentage of observed NES_XY_ ≥ 0, and similarly if NES_AB_ ≤ 0. For the analyses done in this study, the background gene sets used were metabolic pathways defined by the Kyoto Encyclopedia of Genes and Genomes.

### Availability of DGSEA software

R scripts used to 1) calculate DGSEA and GSEA values for two gene sets and 2) generate mountain plots are freely available at: https://github.com/JamesJoly/DGSEA

### Hypoxia Analysis

Gene expression data was downloaded from Ye and Fertig et al. (17) for 31 breast cancer cell lines in 20% oxygen or 1% oxygen. For the results in Fig. 2B-C, the data was median normalized and the log_2_ fold-change was calculated for each gene upon subjection to 1% oxygen to generate ranked gene lists. Then, DGSEA and GSEA were run to generate NES for glycolysis, OxPhos, Hypoxia, and DGSEA. The HALLMARK_HYPOXIA M5891 gene set was used from MSigDB as a benchmark (Supp. Fig. 1). Heatmaps were generated using Morpheus, https://software.broadinstitute.org/morpheus. Mountain plots were generated using. For the results in Fig. 2D, the gene expression data was scaled and centered for all 62 samples and then single-sample GSEA and DGSEA were run for glycolysis, OxPhos, Hypoxia, and DGSEA.

### NCI60 Consumption and Release Analysis

Consumption and secretion rates for glucose and lactate were used from Jain. et al. (19). The raw data was averaged per cell line. Gene expression data was downloaded from Gmeiner et al. (38), then filtered for protein coding transcripts. Gene expression data was centered and scaled across all cell lines, generating rank lists for each cell line. GSEA and DGSEA were run using the weighted Kolmogorov-Smirnov-like statistic to generate normalized enrichment scores (NES) for glycolysis, OxPhos, and DGSEA. The Spearman correlation coefficient was calculated comparing NES to lactate secretion or glucose consumption.

### Cancer Cell Line Encyclopedia Analysis

Gene expression and metabolomics data were downloaded from the CCLE. Gene expression data was centered and scaled across all cell lines, then GSEA and DGSEA were run to generate NES for glycolysis, OxPhos, and DGSEA. The Spearman correlation coefficient was calculated comparing NES to all metabolite abundances. P-values for the Spearman correlation coefficients were adjusted for multiple hypotheses using the Benjamini-Hochberg method.

### Cancer Dependency Map Analysis

Gene expression data was downloaded from the Cancer Cell Line Encyclopedia and was centered and scaled across all cell lines. GSEA and DGSEA were run to generate NES for glycolysis, OxPhos, and DGSEA. Gene essentiality data for each cell line was downloaded from the Cancer Dependency Map (23). The spearman correlation coefficient was calculated between each gene and each NES, generating a rank list reflecting the correlation between gene essentiality and GSEA or DGSEA score. P-values for the Spearman correlation coefficients were adjusted for multiple hypotheses using the Benjamini-Hochberg method. Then, GSEA for all metabolic pathways in the Kyoto Encyclopedia of Genes and Genomes (KEGG) was run on the rank lists. For the results in Fig. 5D, the p-value for over-enrichment was calculated based on the cumulative distribution function of the hypergeometric distribution using: https://systems.crump.ucla.edu/hypergeometric/index.php.

