## Supporting Info for "Differential Gene Set Enrichment Analysis: A statistical approach to quantify the relative enrichment of two gene sets"

Running title: Differential Gene Set Enrichment Analysis

To whom correspondence should be addressed: Nicholas A. Graham, University of Southern California, Los Angeles, 3710 McClintock Ave., RTH 509, Los Angeles, CA 90089. Phone: 213-240-0449;

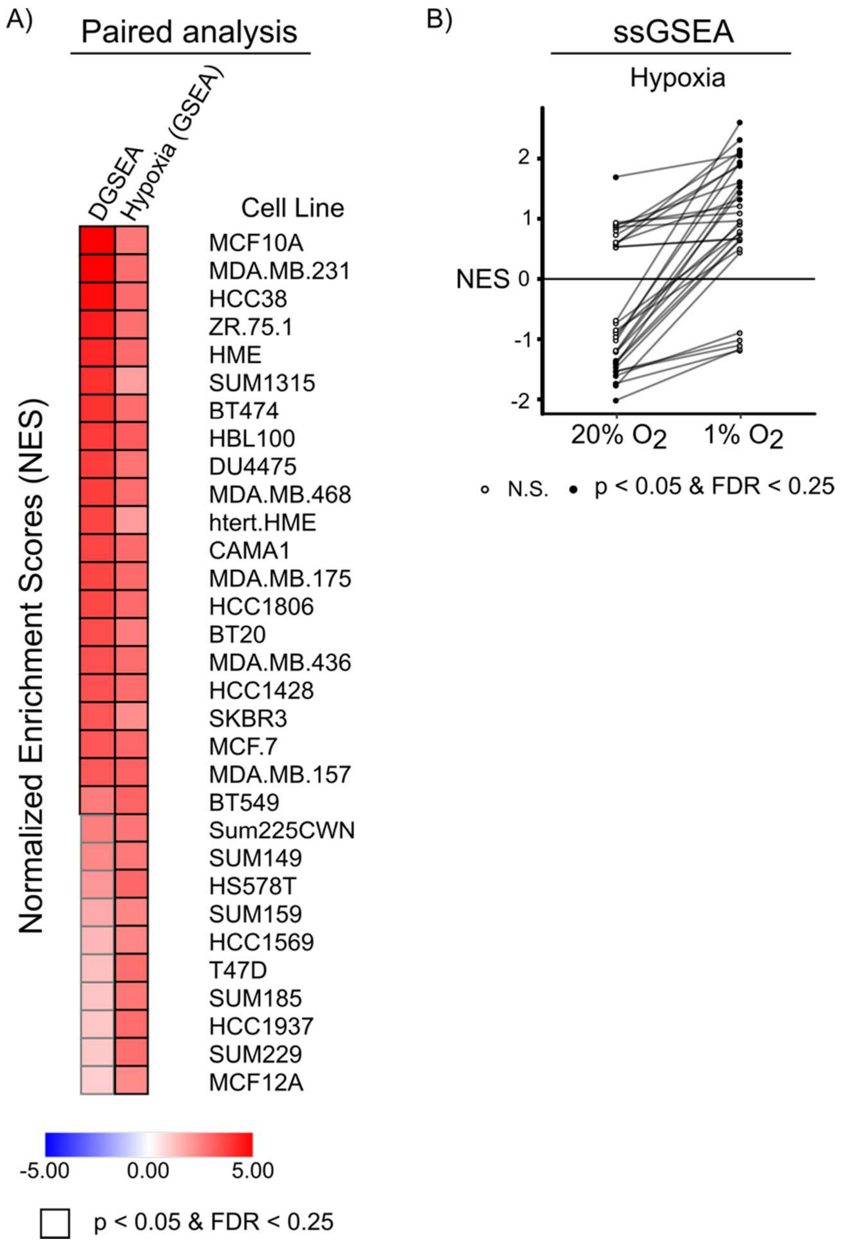


**Supporting Figure 1. Validation of Hypoxia gene signature. A)** To benchmark our analysis in Figure 1, we used a hypoxia gene set (HALLMARKS_HYPOXIA) from the Molecular Signature Database (MSigDB) of the Broad Institute. Paired analysis of hypoxia over normoxia revealed that all breast cancer cell lines upregulated the hypoxia gene signature when subjected to 1% oxygen. **B)** Single sample GSEA (ssGSEA) using the hypoxia gene signature was performed on 62 breast cancer cell lines subjected to either 20% or 1% oxygen. Upon 1% oxygen, most but not all cancer cell lines increased the Normalized Enrichment Score (NES) of the hypoxia gene signature. Filled circles denote nominal p-value < 0.05 and FDR < 0.25. Open circles denote not significant (N.S.).

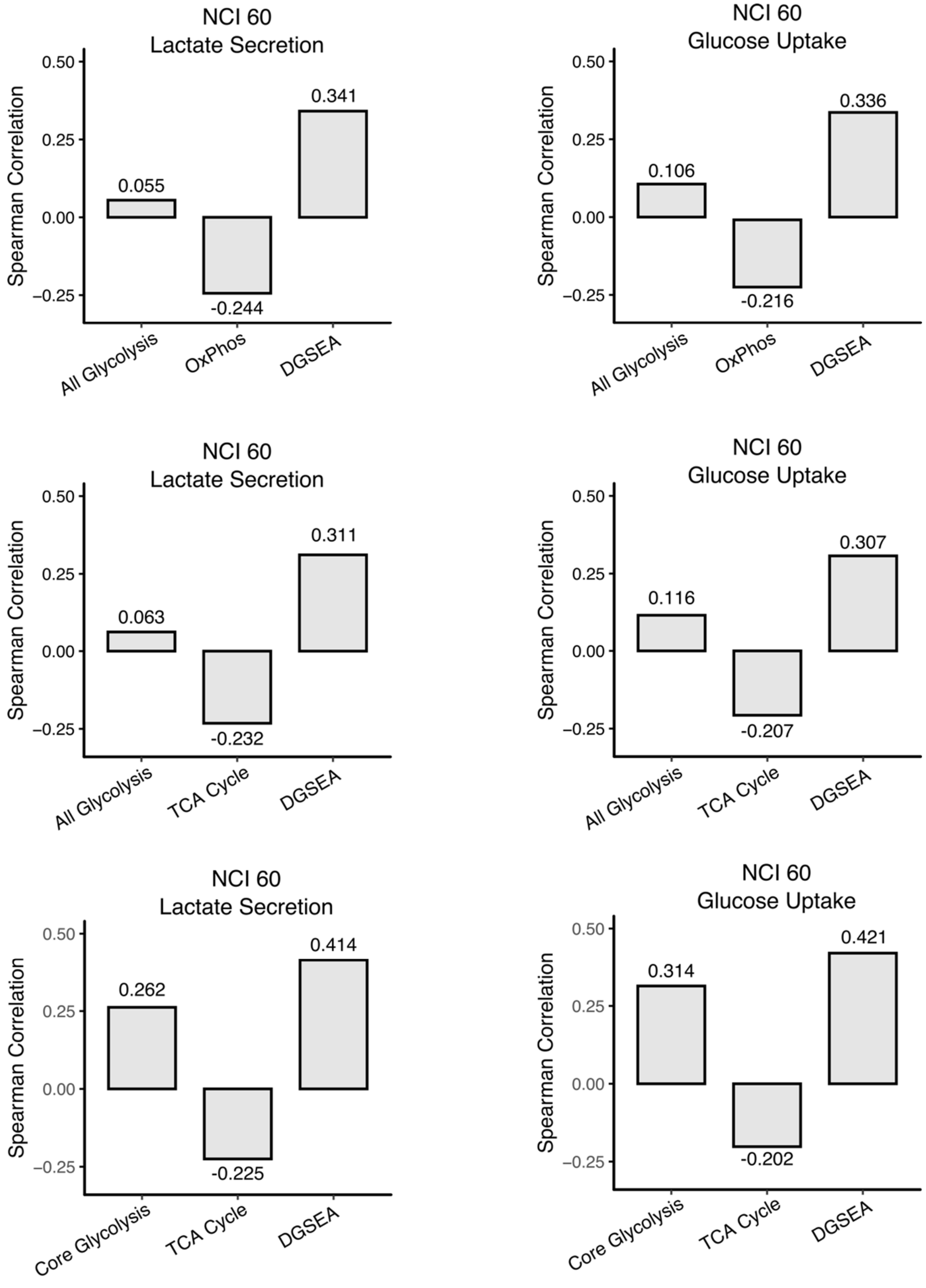


**Supporting Figure 2. Testing Gene sets for DGSEA comparing Glycolysis and Oxidative Phosphorylation.** DGSEA is a better predictor of lactate secretion and glucose uptake than

GSEA regardless of which glycolysis and oxidative phosphorylation gene sets are used. Gene expression data was centered and scaled across 59 of the NCI-60 cancer cell lines and glycolysis, OxPhos, and DGSEA Normalized Enrichment Scores (NES) were calculated for each cell line using the indicated gene sets for glycolysis (All Glycolysis, Core Glycolysis) and oxidative phosphorylation (OxPhos, TCA Cycle). Spearman rank correlation coefficients were calculated between each NES and lactate secretion or glucose uptake data from Jain et al (1).

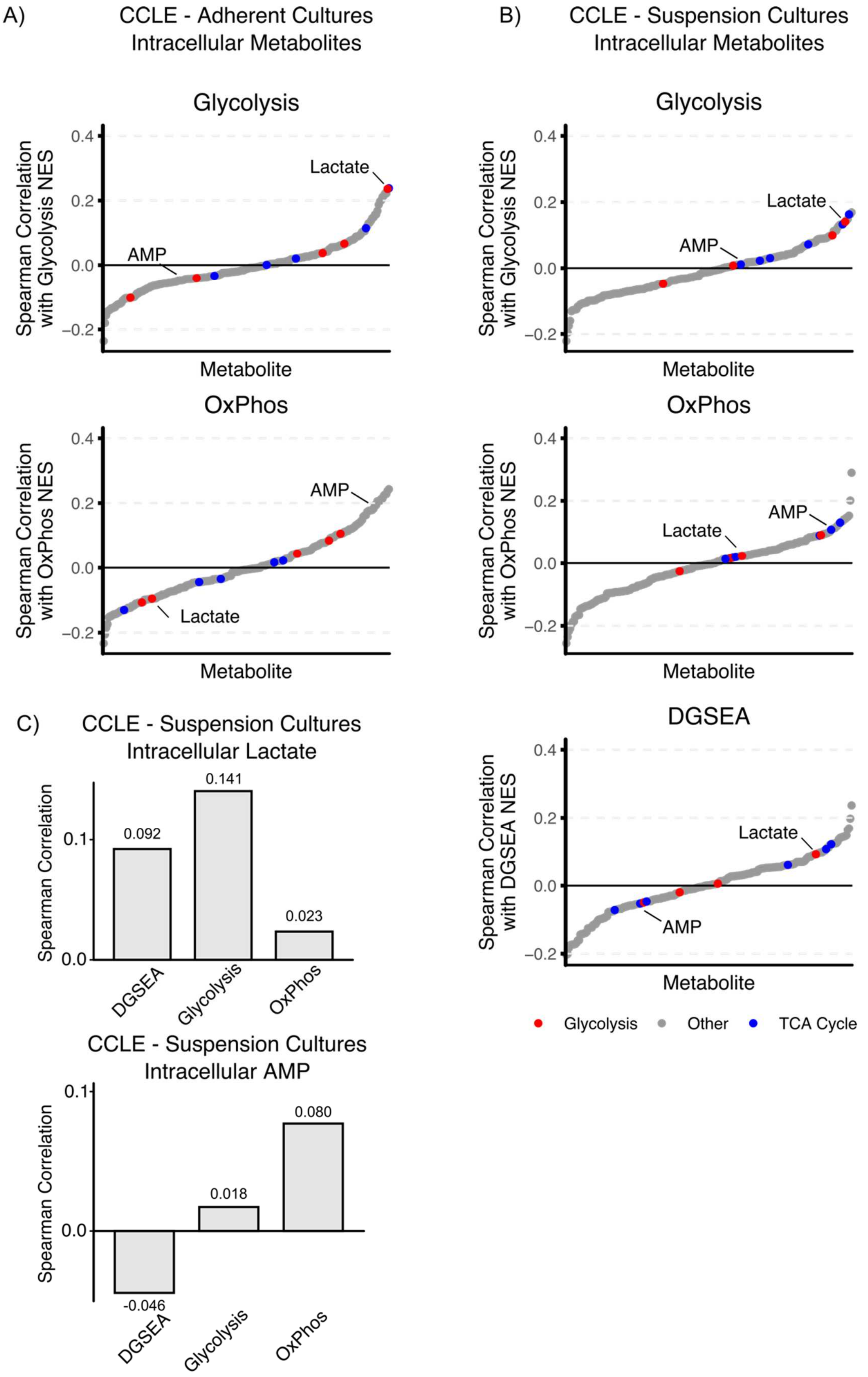
A)
CCLE - Adherent Cultures
Intracellular Metabolites
Glycolysis
Spearman Correlation with Glycolysis NES
Lactate
AMP
Metabolite
OxPhos
Spearman Correlation with OxPhos NES
AMP
Lactate
Metabolite
B)
CCLE - Suspension Cultures
Intracellular Metabolites
Glycolysis
Spearman Correlation with Glycolysis NES
Lactate
AMP
Metabolite
OxPhos
Spearman Correlation with OxPhos NES
AMP
Lactate
Metabolite
DGSEA
Spearman Correlation with DGSEA NES
Lactate
AMP
Metabolite
Glycolysis
Other
TCA Cycle
C)
CCLE - Suspension Cultures
Intracellular Lactate
Spearman Correlation
0.092
0.141
0.023
DGSEA
Glycolysis
OxPhos
CCLE - Suspension Cultures
Intracellular AMP
Spearman Correlation
-0.046
0.018
0.080
DGSEA
Glycolysis
OxPhos

**Supporting Figure 3. DGSEA analysis with steady state metabolite levels in adherent and suspension cells.** RNASeq data was centered and scaled for 836 adherent or 173 suspension cell culture lines in the Cancer Cell Line Encyclopedia. Then the spearman correlation coefficient was calculated for between DGSEA NESs and metabolite abundances. **A)** Lactate was the third most correlated metabolite with glycolysis NES for adherent cancer cell lines. Waterfall plots showing the spearman correlation between all metabolites and glycolysis or OxPhos NES are shown. **B)** Increased lactate and decreased AMP levels did not correlate with DGSEA in cell lines cultured in suspension. Waterfall plots showing the spearman correlation between all metabolites and glycolysis, OxPhos or DGSEA NES are shown. **C)** Individual bar plots showing the correlation between glycolysis, OxPhos, and DGSEA NES and intracellular lactate and AMP for cancer cell lines cultured in suspension are shown.

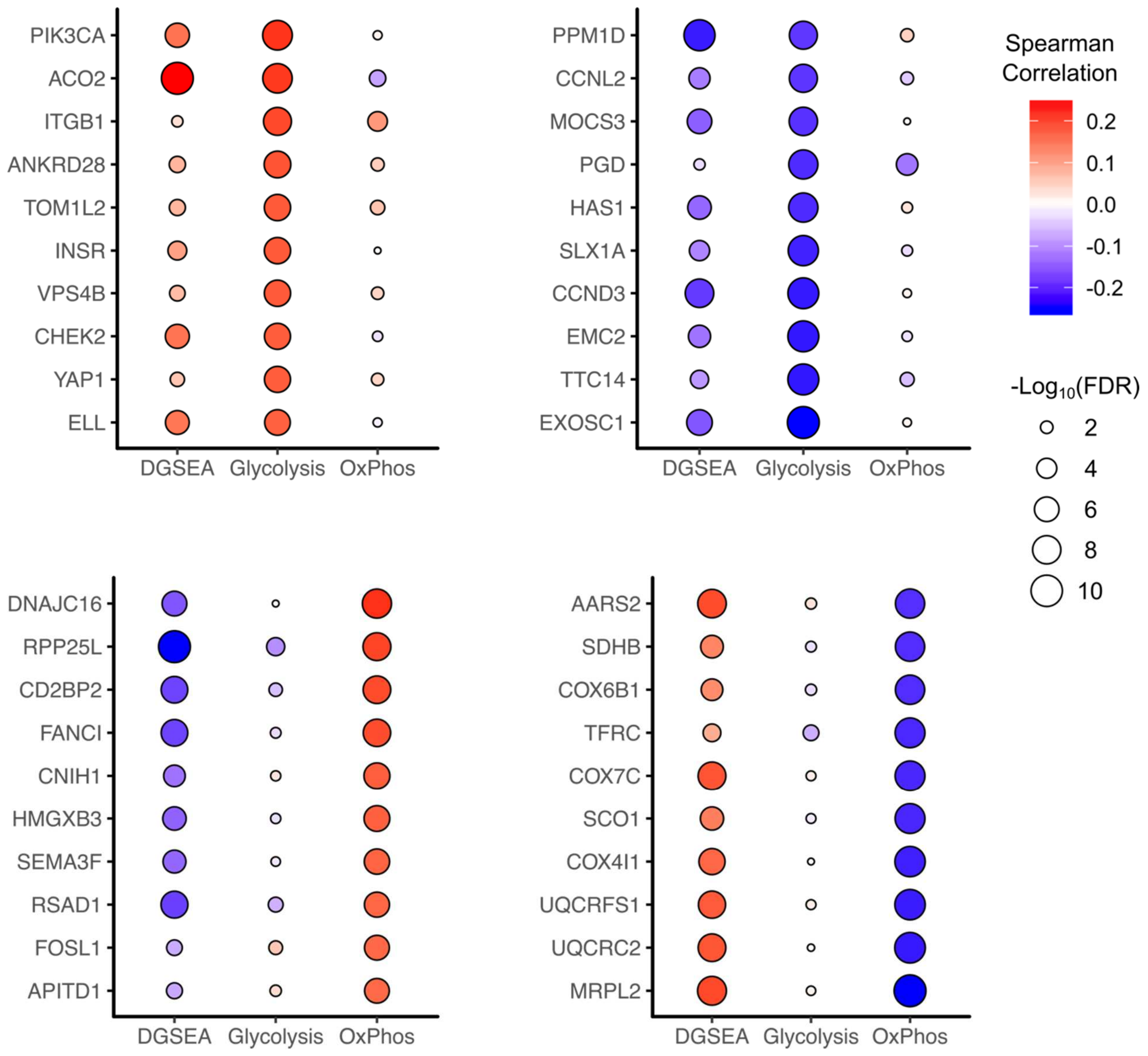


**Supporting Figure 4. Genetic dependencies trend with glycolysis and oxidative phosphorylation.** RNA-Sequencing data was scaled and centered for 1,019 cancer cell lines and DGSEA or GSEA was calculated for each cell line. Spearman correlations were calculated between dependency (CERES score) and the normalized enrichment scores (NES) from DGSEA or GSEA. Spearman correlations for the top 10 and bottom 10 most correlated genes with glycolysis (top) and OxPhos (bottom) are shown. The circle color indicates the correlation coefficient and the circle size indicates the FDR-corrected p-value.

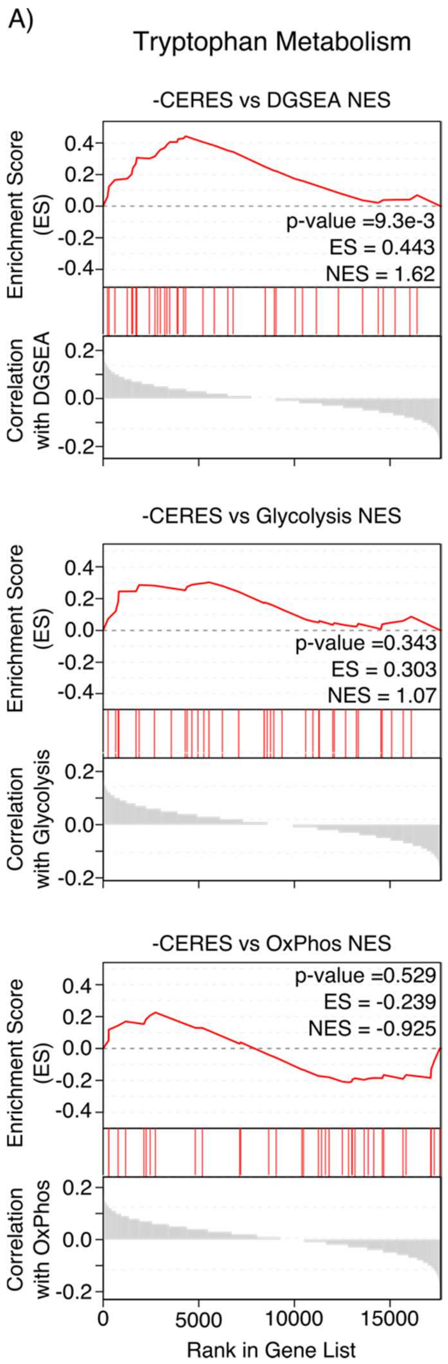


**Supporting Figure 5. Dependency for Tryptophan metabolism increases when glycolysis is high and OxPhos is low in cancer cell lines**. RNA-Sequencing data was scaled and centered for 1,019 cancer cell lines and DGSEA or GSEA was calculated for each cell line. Spearman correlations were calculated between dependency (CERES score) and the normalized enrichment scores (NES) from DGSEA or GSEA. Then, GSEA was run against all KEGG metabolic pathways to identify pathways that exhibit increased or decreased dependency with DGSEA. Mountain plots for Tryptophan Metabolism when ranking the correlation of dependency (-CERES score) with glycolysis, OxPhos, or DGSEA are shown.
